# O Short-branch Microsporidia, Where Art Thou? Identifying diversity hotspots for future sampling

**DOI:** 10.1101/2024.04.01.587430

**Authors:** Megan Gross, Ľubomír Rajter, Frédéric Mahé, David Bass, Cédric Berney, Nicolas Henry, Colomban de Vargas, Micah Dunthorn

**Affiliations:** Natural History Museum, University of Oslo, 0562 Oslo, Norway; Department of Ecology, University of Kaiserslautern-Landau RPTU, 67663 Kaiserslautern, Germany; Institute for Zoology, University of Cologne, 50923 Cologne, Germany; CIRAD, UMR PHIM, 34398 Montpellier, France; PHIM, Univ Montpellier, CIRAD, INRAE, Institut Agro, IRD, 34398 Montpellier, France; Cefas, International Centre for Aquatic Animal Health, Weymouth, Dorset DT4 8UB, United Kingdom; Sustainable Aquaculture Futures, Biosciences, College of Life and Environmental Sciences, University of Exeter, Stocker Road, Exeter EX4 4QD, United Kingdom; Department of Life Sciences, The Natural History Museum, London SW7 5BD, United Kingdom; CNRS, Sorbonne Université, FR2424, ABiMS, Station Biologique de Roscoff, 29680 Roscoff, France; Sorbonne Université, CNRS, Station Biologique de Roscoff, UMR7144, ECOMAP, 29680 Roscoff, France; Research Federation for the study of Global Ocean Systems Ecology and Evolution, FR2022/Tara GOSEE, 75016 Paris, France

**Keywords:** metabarcoding, parasites, protists, SSU-rRNA, SSU-V4

## Abstract

Short-branch Microsporidia were previously shown to form a basal grade within the expanded Microsporidia clade and to branch near the classical, long-branch Microsporidia. Although they share simpler versions of some morphological characteristics, they do not show accelerated evolutionary rates, making them ideal candidates to study the evolutionary trajectories that has led to long-branch microsporidians unique characteristics. However, most sequences assigned to the short-branch Microsporidia are undescribed, novel environmental lineages for which the identification requires knowledge of where they can be found. To direct future isolation, we used the EukBank database of the global UniEuk initiative that contains the majority of the publicly available environmental V4 SSU-rRNA gene sequences of protists. The curated OTU table and corresponding metadata were used to evaluate the occurrence of short-branch Microsporidia across freshwater, hypersaline, marine benthic, marine pelagic, and terrestrial environments. Presence-absence analyses infer that short-branch Microsporidia are most abundant in freshwater and terrestrial environments, and alpha- and beta-diversity measures indicate that focusing our sampling effort on these two environments would cover a large part of their overall diversity. These results can be used to coordinate future isolation and sampling campaigns to better understand the mysterious evolution of microsporidians unique characteristics.

## 1. Introduction

Parasites comprise a large fraction of Earth’s biodiversity (Dobson et al., 2008; Loker and Hofkin, 2022; Mahé et al., 2017) and their transition from free-living life forms to a parasitic lifestyle occurred in numerous independent events. This convergent evolution among phylogenetically unrelated taxa resulted in strategies of host-invasion, transmission between, and survival within their hosts (Poulin, 2011; Poulin and Randhawa, 2015). One clade of obligate intracellular parasites is the Microsporidia (Keeling and Fast, 2002; Vávra and Lukeš, 2013; Weiss and Becnel, 2014). While the diversity of Microsporidia and their infection mechanisms are somewhat understood, it is still not clear how their unique characteristics, and thus how this clade itself, evolved (Keeling and Fast, 2002).

Microsporidians infect a wide range of animals and some protists (Adl et al., 2019; Becnel and Andreadis, 1999; Foissner and Foissner, 1995; Vávra and Lukeš, 2013; Vossbrinck and Debrunner-Vossbrinck, 2005). Thus far, more than 1,300 species are described (Franzen, 2008; Weiss and Becnel, 2014), many of them being harmful and emergent pathogens of socio-economic importance (Fries, 1993; Keeling and Fast, 2002; Kent et al., 1989; Stentiford et al., 2016; Weber et al., 1994). While microsporidians display a wide complexity in their infection pathways using a unique polar filament for host invasion, they otherwise have had a rather reductive evolution in their cellular organization (Dean et al., 2018; Katinka et al., 2001; Keeling and Corradi, 2011). The loss of many DNA repair enzymes (Gill and Fast, 2007) and the presence of many fast-evolving genes (Thomarat et al., 2004), may have enhanced the process of genome reduction thereby playing a major role in the evolution of Microsporidia (Cuomo et al., 2012).

Accelerated evolutionary rates caused long-branch attraction artefacts and initial phylogenetic inferences placed the microsporidians as an early branching eukaryote clade (Keeling and McFadden, 1998; Vossbrinck et al., 1987). Subsequent phylogenetic analyses that took into account variable base substitutions and included numerous loci changed our understanding of their placement within the eukaryotic tree of life (Corradi and Keeling, 2009; Park and Poulin, 2021). It is now widely accepted that the microsporidians are closely related to fungi (Brown and Doolittle, 1999; Corradi and Keeling, 2009; Gill and Fast, 2006; Hirt et al., 1999; Keeling, 2014, 2003; Strassert and Monaghan, 2022; Voigt et al., 2021). However, although many studies made important contributions to unravel their diversity and phylogeny, as well as shedding light on the unique characteristics that make the microsporidians such pivotal parasites, it still remains unclear how these characteristics evolved.

Bass et al. (2018) expanded our understanding of the relationship between microsporidia and many different microsporidian-like protists that were previously assumed to group together with rozellids (parasites of Chytridiomycetes, Blastocladiomycetes and Oomycetes) within the ‘cryptomycotan’ clade. They used small subunit rRNA (SSU-rRNA) from sequenced isolates and environmental metabarcoding data to infer that these microsporidian-like lineages form a basal grade of parasites that group together with metchnikovellids and classical microsporidia, thereby demonstrating that the phylogenetic scope of the microsporidians is greater than originally assumed. This expanded microsporidian clade includes all classical Microsporidia, Metchnikovellida and Chytridiopsida, which they collectively named ‘long-branch Microsporidia’ (referring to the relatively long branches within phylogenetic trees), and the basal grade with less divergent SSU-rRNA gene sequences, which they named ‘short-branch Microsporidia’.

The short-branch Microsporidia include the partially-characterized lineages *Mitosporidium, Morellospora*, *Nucleophaga*, and *Paramicrosporidium*, which are variously known to be parasites of *Daphnia* (Haag et al., 2014) and different amoebae (Corsaro et al., 2020, 2014; Michel et al., 2009b, 2000). They share some morphological traits with the long-branch Microsporidia, such as simpler versions of the polar filament (with a similar function) but differ greatly in other characteristics. For example, they do not show rapid rates of evolution and have less reduced genomes that are more similar to those of *Rozella* and canonical Fungi (Haag et al., 2014; Quandt et al., 2017). The short-branch Microsporidia also include many uncharacterized environmental lineages that may be parasitic as well (Doliwa et al., 2021). By further investigating the short-branch Microsporidia, we can better understand the intriguing evolution that occurred among the classical long-branch Microsporidia, which ultimately resulted in trait reductions and increased complexity of their extrusion apparatus. However, as the short-branch Microsporidia are undersampled and understudied, we don’t yet know where to focus attempts in order to isolate more of them, to further investigate the partially-characterized lineages and to newly characterize the environmental lineages.

In this study, we used data from Berney et al. (2023) that collected and analyzed all available environmental metabarcoding data of the protistan hypervariable V4 region of the SSU-rRNA gene. After extracting short-branch Microsporidia operational taxonomic units (OTUs), we asked in which environments short-branch Microsporidia are present and if there are differences in their abundance and diversity throughout these environments. Our results pave the road for future studies in which we aim to better understand the evolution of Microsporidia, by directing where we should go to isolate and sequence more short-branch Microsporidia.

## 2. Results

### 2.1 Sampling and sequencing

Of the 13,040 samples taken from across the globe, the majority came from marine pelagic environments (62.89%), followed by terrestrial (19.36%), marine benthos (13.58%), freshwater (3.92%), and hypersaline (0.25%) environments (**Figure 1**). The majority of the terrestrial samples were taken in areas in the northern hemisphere, especially within Europe. There were also many countries for which no samples targeting the V4 region were available including the whole African continent, Australia and also parts of South America such as Brazil. From the initial dataset, 1,403,019,176 environmental V4 sequencing reads clustered into 460,147 OTUS that were taxonomically assigned to the protists. After filtering out all non-short-branch microsporidian sequences, 6,796,304 (0.48%) reads remained that were clustered into 1,741 short-branch microsporidian OTUs. From these short-branch Microsporidia data, 99.2% of the reads and 98.4% of the OTUs were more than 85% similar to references in the EukRibo (Berney et al., 2022) reference database (**Figure 2**).

**Figure 1:**
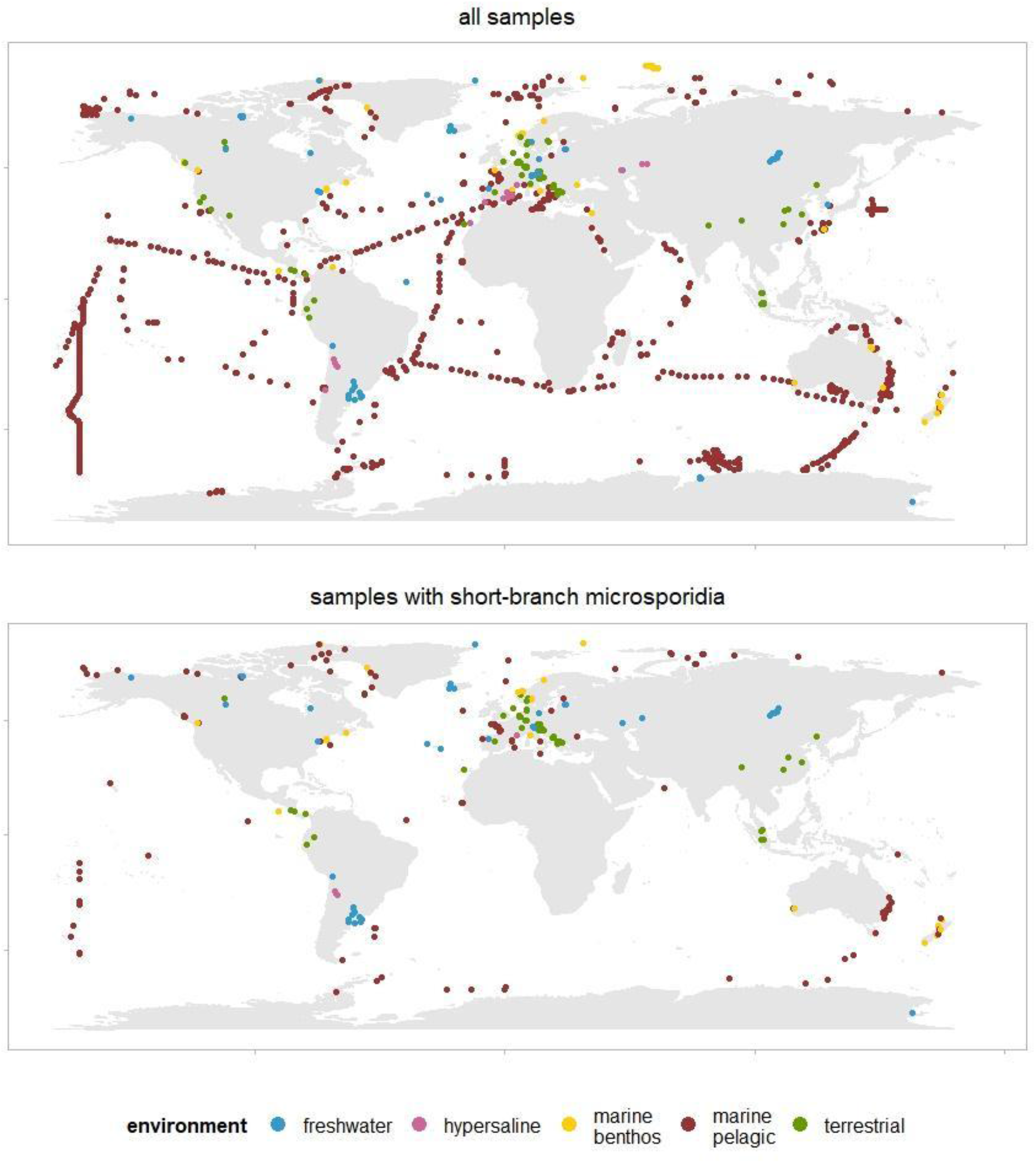
Map of all sample locations, and map of samples that include short-branch microsporidia. Colours represent the environment in which the samples were collected.

**Figure 2:**
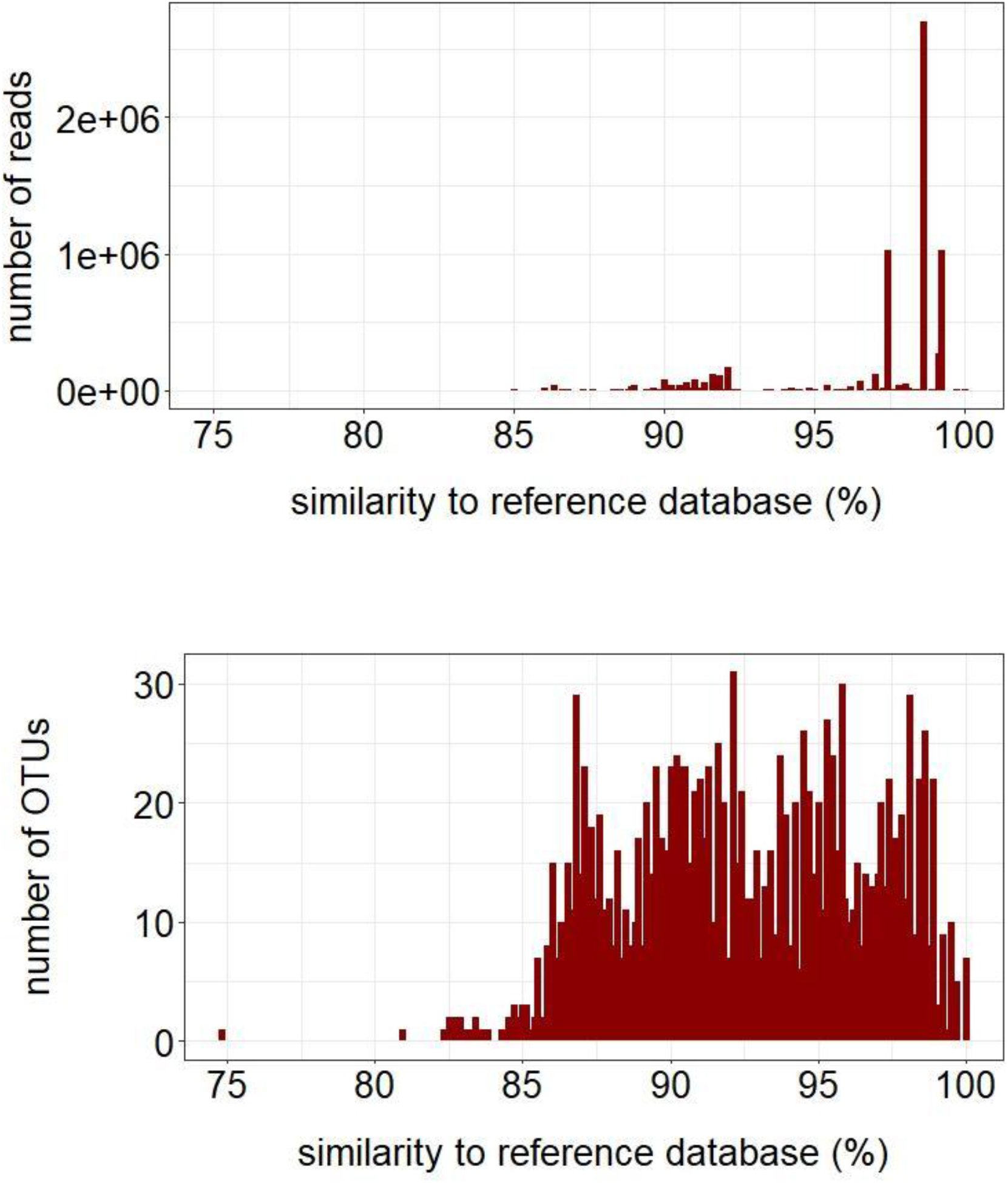
Similarity of short-branch Microsporidia to the taxonomic reference database for a) number of reads and b) number of OTUs.

### 2.2 Distributions across environments

Of the 13,040 samples, we found short-branch microsporidian OTUs within 3,463 (26.55%) samples (**Figure 1**). A majority of these samples (1,818) and OTUs (1,279) were found in terrestrial environments (**Figure 3**). Although there were more samples from marine environments, freshwater samples accounted for 879 OTUs compared to 584 for marine pelagic and 567 OTUs for marine benthos. A similar picture was found when searching for unique and shared OTUs across the different environments. Terrestrial environments had the highest number of unique OTUs (603) followed by freshwater (191). There were 151 OTUs shared between freshwater and terrestrial environments, 219 that occurred within every environment with the exception of hypersaline, and three OTUs occurred in all five environments. These three OTUs found in all five environments had a high similarity to accessions in the taxonomic reference database (>98%). Two of the OTUs were 98.26% similar to each other and had *Paramicrosporidium clone LKM-46* (van Hannen et al., 1999) as their closest reference. The third OTU was 82.48% and 82.78% similar to the other two and had *Paramicrosporidium clone MPE1-23* as the closest reference (Nakai et al., 2012).

**Figure 3:**
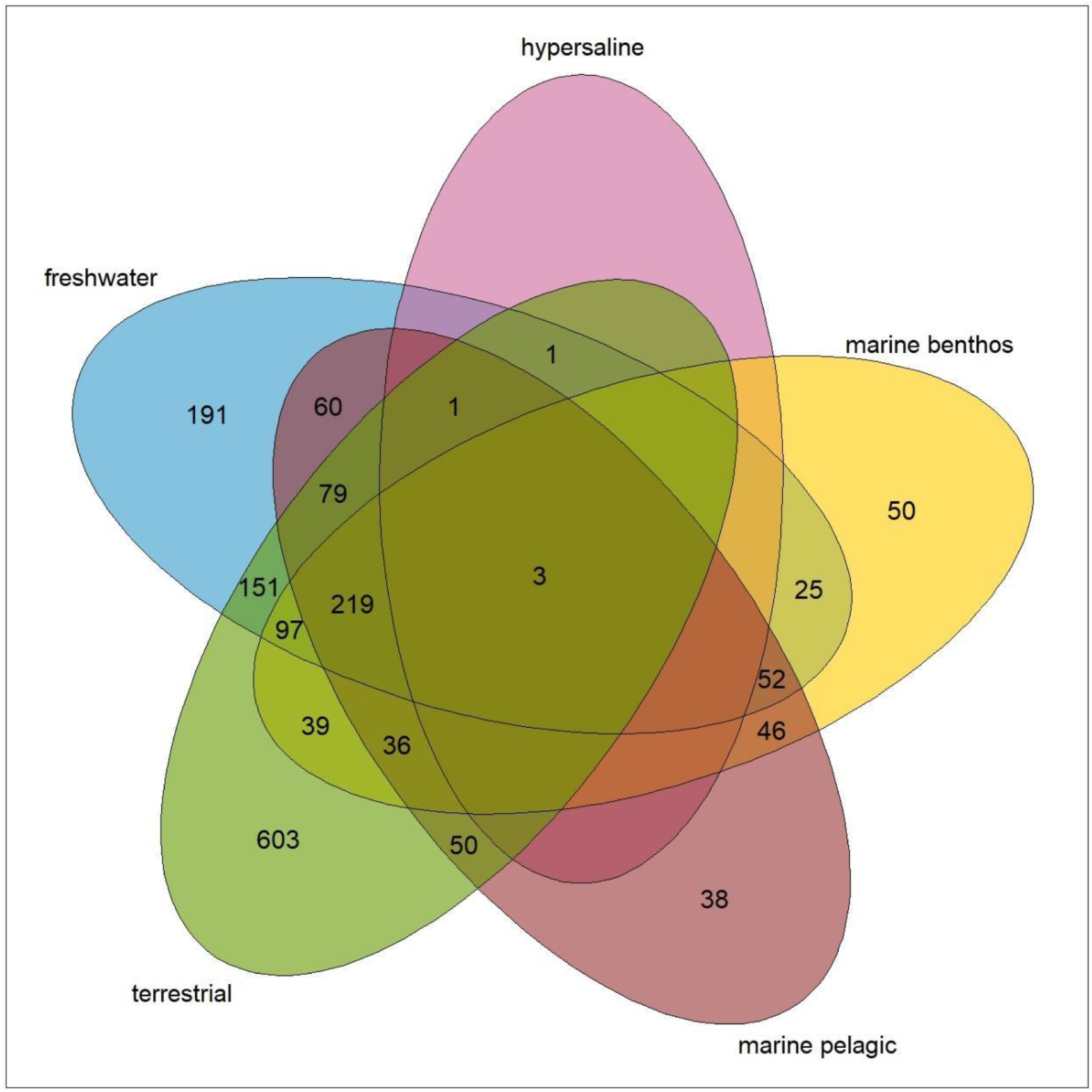
Venn-diagramm showing the number of unique and shared OTUs across the five investigated environments. Sections in which no number appears indicate zero OTUs.

### 2.3 Community richness and heterogeneity

Observed richness of short-branch microsporidian OTUs showed that the average number of OTUs per sample was highest for terrestrial, followed by freshwater environments (**Figure 4**). Estimated richness using Chao1 predicted slightly higher diversity values for the different environments. Terrestrial had the highest diversity (Chao1: 14 ± 18.82 SE) followed by freshwater (Chao1: 9.75 ± 43.20 SE). Marine benthos, marine pelagic and hypersaline showed very low average diversity estimates (respectively 1 ± 26.97 SE, 1 ± 13.80 SE, 1 ± 1.22 SE) and differed significantly from freshwater and terrestrial (Suppl. Table 1). Although most samples showed an observed richness of <50 OTUs per sample, 11 freshwater and six marine benthos samples were found with >100 OTUs per sample. Among these, three out of 11 freshwater samples sites were from lake Pollevann in Norway (unpublished study), seven came from lake Sanabria in Spain (unpublished study), and one sample originated from lake Augstsee in Austria. All of the six marine benthos samples were from Norwegian fjords. In contrast to the differences in observed and estimated OTU richnesses, there were no clear differences in the communities of short-branch Microsporidia across the environments (**Figure 5**). Non-metric multidimensional scaling showed that the majority of samples clustered together with the exception of some marine benthic, marine pelagic, and terrestrial samples.

**Figure 4:**
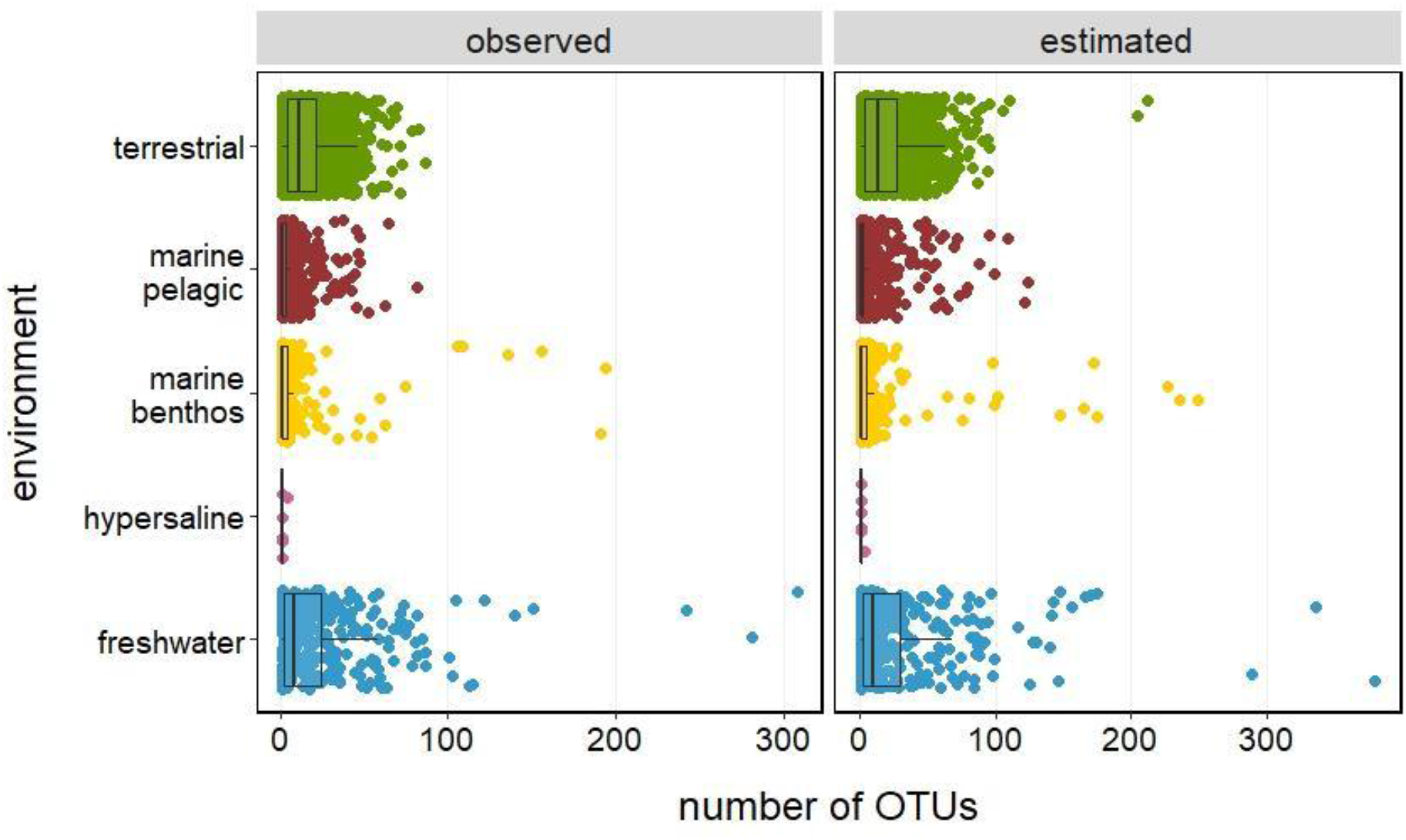
Alpha-diversity per environment for observed number of OTUs and estimated number of OTUs using Chao1.

**Figure 5:**
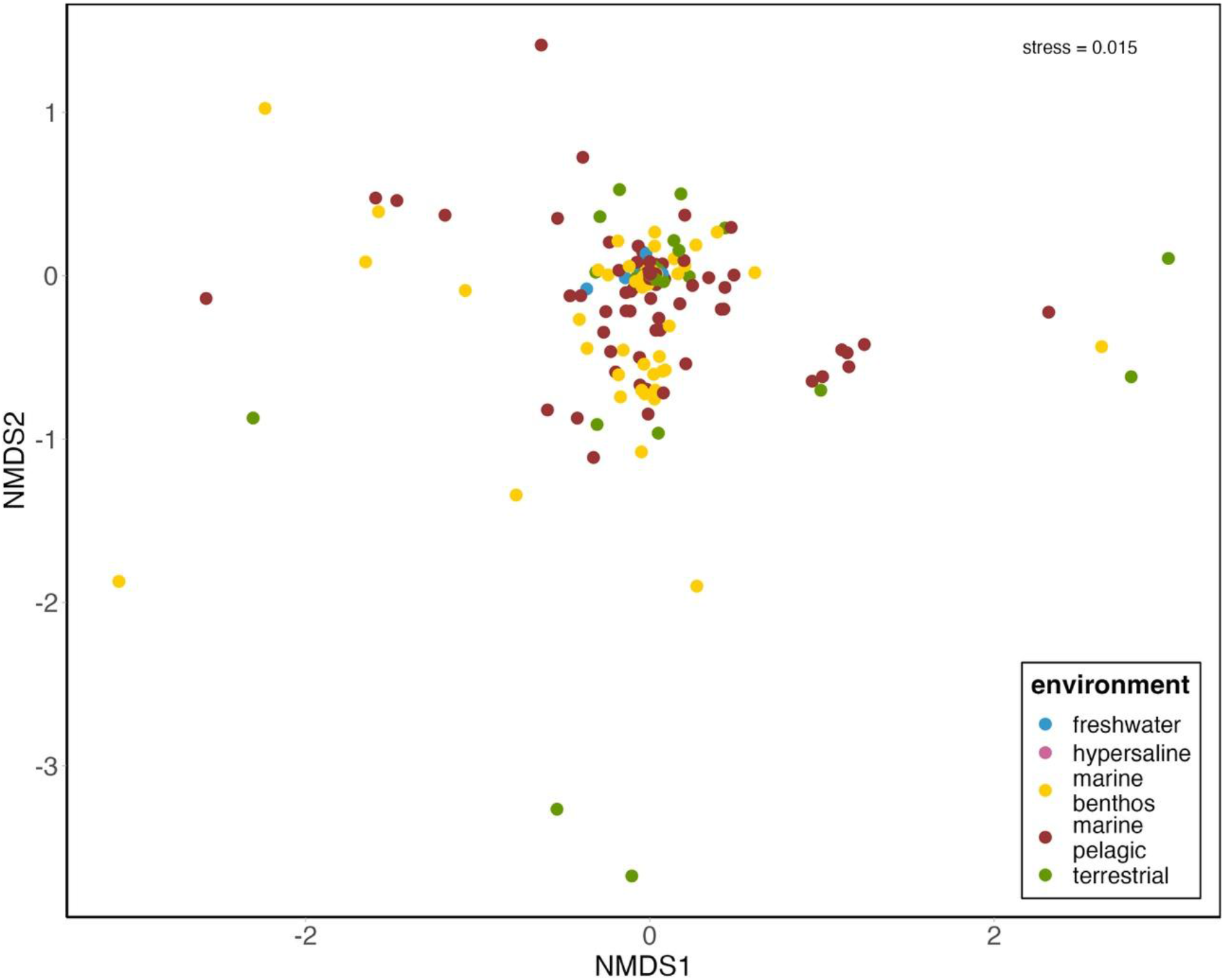
Non-metric multidimensional scaling plot of all samples across the different environments (Dissimilarity measure = Jaccard-Index).

## 3. Discussion

Our data show that short-branch Microsporidia are ubiquitous protists that occur in all of the investigated environments with many of them not closely related to already sequenced isolates (**Figure 1, 2**). Although widespread, we nevertheless found differences in their abundance across environments, with terrestrial accounting for the highest number of samples with short-branch Microsporidia, followed by freshwater (**Figures 1, 4**). However, since there were no clear community differences between environments (**Figure 5**), we can infer that sampling in terrestrial and freshwater environments would cover a large fraction of the overall diversity of the short-branch Microsporidia. These findings can be used to direct future isolation of short-branch Microsporidia.

Using environmental sequencing data targeting the SSU-rRNA has not only revealed an incredible diversity of unicellular eukaryotes (Logares et al., 2020; Massana et al., 2015; Santoferrara et al., 2020; de Vargas et al., 2015), but also advanced our understanding of microsporidian diversity over the last few decades. Many of these studies led to some novel findings, and highlighted that Microsporidia are widespread parasites with a large part of their diversity that remains undescribed (Ardila-Garcia et al., 2013; Dubuffet et al., 2021; Murareanu et al., 2021; Williams et al., 2018). However, since short-branch Microsporidia are generally referred to as ‘microsporidian-like’, and because they are frequently considered to group indistinguishably from the rozellids, published studies often did not include them when investigating the diversity and distribution of long-branch Microsporidia. Additionally, it was not until Bass et al. (2018) that the relationship between long-branch and short-branch Microsporidia was solidly known. Here, with our dataset of 3,463 samples and only selecting short-branch microsporidian sequences, our results show that this high diversity not only applies to the long-branch Microsporidia, but also applies to the short-branch Microsporidia (**Figure 1**). We also found that around 30% of the sequences analyzed here had no close reference (less than 90% similarity) within the taxonomic reference database (**Figure 2**), and may therefore represent further undescribed lineages, underpinning the need to isolate and sequence more short-branch Microsporidia.

Beyond the standard problems of using an environmental metabarcoding approach to evaluate protistan environmental diversity (Santoferrara et al., 2020), the uneven sampling in our study could have led to an underestimation of short-branch Microsporidia in some parts of Earth. In particular, large parts of Africa, Australia, North and South America are underrepresented or not represented at all in this study. Even with these limitations in mind, our results still show that terrestrial and freshwater environments harbor a large diversity of short-branch Microsporidia that warrant further exploration.

We found that 219 of the short-branch microsporidian OTUs were shared between freshwater, marine benthos, marine pelagic and terrestrial environments, and 151 OTUs were shared between freshwater and terrestrial environments, thus increasing our level of confidence in the reality of this diversity (**Figure 3**). Although early classifications grouped Microsporidia into the three groups ‘Aquasporidia’, ‘Marinosporidia’ and ‘Terresporidia’ (Vossbrinck and Debrunner-Vossbrinck, 2005), more recent studies have shown that many species are associated with more than one environment (Murareanu et al., 2021). This heterogeneity may be the result of their hosts being widespread and occurring in different environments themselves (Park et al., 2020). For example, Murareanu et al. (2021) found that 27.4% of microsporidian hosts were classified to occur in more than one environment. Furthermore, some microsporidians have been classified as generalists and can infect more than one host species of different environments (Stentiford et al., 2016). Both of these scenarios could apply to many lineages of short-branch Microsporidia, especially for those three OTUs that were associates with all environments. Despite the many shared OTUs, we also found a considerable number of unique OTUs within freshwater and terrestrial environments (**Figure 3**; 191 and 603 OTUs, respectively) that may allow to resolve some of the uncharacterized diversity. In addition, although the large diversity of environmental lineages of short-branch Microsporidia was first shown in neotropical rainforest soils by Bass et al. (2018), we found more diversity within 17 samples taken from freshwater and marine benthos environments of the northern hemisphere. These samples came from the freshwater lake Pollevann and fjords close to Oslo, Norway as well as from lake Sanabria, Spain.

The greater diversity and abundance of short-branch Microsporidia in freshwater lakes is also in agreement with the assumption that evolutionary transitions to long-branch Microsporidia have occurred in freshwater environments due to the accessibility to free-living amoeba as transfer hosts (Corsaro et al., 2019). However, our results also showed that many short-branch microsporidian OTUs were found within samples from Norwegian fjords. One possible explanation may be that many of the free-living amoeba that have been identified as hosts have also been described from marine environments (Page, 1977), or related amoebae in marine and non-marine environments are similarly infected by short-branch Microsporidia. This could also apply to the three OTUs that were shared across all five investigated environments and who had *Paramicrosporidium* as the closest genus. These references were originally found in freshwater environments. The original sequence assigned to *Paramicrosporidium clone MPE1-23* was isolated from aquatic moss pillars from lake Hotoke-Ike in Skarvsnes, Antarctica (Nakai et al., 2012). The original sequence assigned to *Paramicrosporidium clone LKM-46* was isolated from cultures containing water from lake Ketelmeer, Netherlands and various sources of detritus (van Hannen et al., 1999). However, it was not clear in those original studies that the sequences were derived from parasitic species and who their potential hosts were.

The findings of this study also further our understanding about what is known about the partially characterized lineages so far. For example, *Mitosporidium daphniae* infects the water flea *Daphnia* that occurs in freshwater habitats and *Nucleophaga terricolae* infects free-living amoebae that were isolated from the bark of trees (Corsaro et al., 2016; Michel et al., 2012). The same applies to other partially-characterized lineages that were also isolated from hosts derived from freshwater or terrestrial samples (Corsaro et al., 2020, 2016; Michel et al., 2009a, 2000). A recent study investigating host-parasite interactions in a freshwater lake found that phytoplankton and microzooplankton (e.g., ciliates) are potential hosts of Microsporidia and suggested that they could be of greater importance for the functioning of lake ecosystems than previously known (Chauvet et al., 2022). The same could apply to the short-branch Microsporidia, since many of their identified hosts are protists and other small animals.

## 4. Conclusion

Using environmental metabarcoding data from samples taken across the world, our study shows that there is a tremendous diversity of short-branch Microsporidia, especially in freshwater and terrestrial environments. In specific, some samples originating from Norway and Spain were particularly rich in short-branch Microsporidia. These data can be used to direct future sampling campaigns, with the goal of isolating and further characterizing partially-described as well as novel-environmental lineages. Furthering our knowledge of these lineages of short-branch Microsporidia may allow us to better understand the evolution that occurred among the long-branch Microsporidia.

## 5. Methods

### 5.1 Dataset

Microsporidian sequences and their corresponding metadata used in this study came from a beta version of the EukBank database (Berney et al., 2023) as part of the global UniEuk initiative (Berney et al., 2017). Briefly, metadata from each bioproject was downloaded from public repositories such as European Nucleotide Archive (EMBL/EBI-ENA) and the Sequence Read Archive (NCBI). Environmental V4 SSU-rRNA raw sequences were downloaded from the EMBL/EBI-ENA EukBank umbrella project, clustered with Swarm v3 using default parameters with the fastidious option on (Mahé et al., 2022), checked for chimeras using VSEARCH (Rognes et al., 2016), merged with MUMU (Mahé 2021), and taxonomically assigned using the stampa pipeline (https://github.con/frederic-mahe/stampa/) and EukRibo v1 reference database (2020-07-16). From this, an OTU (operational taxonomic unit) occurrence table was generated. OTU-, taxonomy- and metadata-table can be found in the supplements (Suppl. File 1-3). All further steps were conducted using R v4.3.1. The codes are available in HTML format (Suppl. File 4).

### 5.2 Modification of OTU Table

Modifications of the dataset were performed using the R package tidyverse v1.3.1 (Wickham et al., 2019). The OTU-, taxonomy- and metadata-tables were matched against each other to make sure that OTUs and samples are identical. To verify that the dataset only contained OTUs annotated to the short-branch Microsporidia the taxonomy table was searched for the key words ‘canonical’ and ‘classical’ Microsporidia. All OTUs to which this applied were removed. To correctly infer the environment in which short-branch microsporidia can be found, every sample was assigned to either of the five different environments i.e. ‘freshwater’, ‘hypersaline’, ‘marine benthic’, ‘marine pelagic’ or ‘terrestrial’ using the information provided in the metadata table. However, for some samples there was not enough information provided in the metadata table itself to correctly assign it into one of the environments. To address this knowledge gap, we searched inside the EMBL-EBI Ontology Code for equivalent information and added it to the metadata table.

### 5.3 Statistical analysis

Analyses and visualizations were performed with the R packages tidyverse v1.3.1 (Wickham et al., 2019), ggpubr v0.4.0 (Kassambara, 2020), rstatix v0.7.0 (Kassambara, 2021), maps v3.4.0 (Brownrigg, 2021), phyloseq v1.34.0 (McMurdie et al., 2020), metagMisc v0.0.4 (Mikryukov, 2021), venn v1.10-5 (Dusa, 2021). All analyses were performed on the unrarefied dataset to ensure that samples are not lost due to poor sample coverage. However, as the sequencing depth varies between different studies due to different sampling efforts and procedures, all analyses were tested on the rarefied dataset as well, to make sure that this did not influence the outcome. The quality of the taxonomic annotation was examined based on the number of OTUs per similarity value to the reference database and was visualized as bar charts. Sampling maps, based on latitude/ longitude information, were created to compare the total amount of samples against samples for which short-branch microsporidian OTUs were found. Samples were color-coded based on the five different environments. To compare the number of unique and shared OTUs present in the different environments, Venn diagrams were created. Additionally, OTUs occurring in all investigated environments were aligned with MUSCLE v3.8.425 (Edgar, 2004) to check for sequence similarity. For community analysis, OTU-, metadata- and taxonomy-table were transformed and then merged into a phyloseq object. Observed and estimated richness (Chao1) was compared between the different environments and results were visualized as boxplots. Wilcoxon rank sum tests were used to test for pairwise differences in richness among the environments (we recognized the dataset was not normally distributed because many samples were low abundant, but we retained all samples to get a complete view of the distributions across the environments). To investigate similarity patterns between the five different environments the OTU table was transformed into presence-absence data and a non-metric multidimensional scaling (NMDS), using the Jaccard metric, was performed.

## Supporting information

OTU table

Taxonomy of OTUs

Metadata of OTUs

Computational codes

Wilcoxon rank sum test for pairwise differences in richness among the environments

## Acknowledgements

This work was funded by an Erasmus+ grant to M.G.

## Notes

### Competing Interest Statement

The authors have declared no competing interest.

