## Supplementary material for "O Short-branch Microsporidia, Where Art Thou? Identifying diversity hotspots for future sampling": Computational codes

### SB-microsporidia-codes

Megan Gross

04/01/2024

#Input files: ASV-Table: eukbank_18SV4_asv.table Metadata-Table: eukbank_18SV4_asv.metadata Taxonomy-Table: eukbank_18SV4_asv.taxo

#### LIBRARIES

#### DATASET

##### Separate datasets based on envrionments

```
land <-
  filter(metadata_table,
         envplot %in% c("land_soil", "land_sediment", "land_org"))
freshwater <-
  filter(metadata_table,
         envplot %in% c("land_freshwater", "land_water"))
marine <-
  filter(metadata_table,
         envplot %in% c("marine_sediment", "marine_org", "marine_water"))
unknown <- filter(metadata_table, envplot %in% c(""))
marine_ice <- filter(metadata_table, envplot %in% c("marine_ice"))
```

```
marine_pelagic <- metadata_table %>%
  filter((material == "sea ice (ENVO_00002200)") |
           (biome == "estuarine biome (ENVO_01000020)") |
           (
             biome == "marine biome (ENVO_00000447)"  &
               material == "algal material (ENVO_01001189)"
           ) |
           (
             biome == "marine biome (ENVO_00000447)"  &
               material == 'sea water (ENVO_00002149)'
           ) |
           (
             biome == "marine benthic biome (ENVO_01000024)"  &
               material == "sea water (ENVO_00002149)"
           ) |
           (
             biome == "marine neritic benthic zone biome (ENVO_01000025)"  &
```

```
              material == "sea water (ENVO_00002149)"
            ) |
            (
              biome == "marine pelagic biome (ENVO_01000023)"  &
                material == "sea water (ENVO_00002149)"
            ) |
            (
              biome == "marine pelagic biome (ENVO_01000023)"  &
                material == "organismal entity (PCO_0000031)"
            ) |
            (
              biome == "oceanic epipelagic zone biome (ENVO_01000035)"  &
                material == "sea water (ENVO_00002149)"
            ) |
            (
              biome == "marine benthic biome (ENVO_01000024)"  &
                material == "sea water (ENVO_00002149)"
            ) |
            (
              biome == "oceanic epipelagic zone biome (ENVO_01000035)"  &
                material == "water (ENVO_00002006)"
            )
)


marine_benthos <- metadata_table %>%
  filter((biome == "marine abyssal zone biome (ENVO_01000027)") |
           (biome == "marine benthic biome (ENVO_01000024)") |
           (
             biome == "marine biome (ENVO_00000447)"  &
               material == "organismal entity (PCO_0000031)"
           ) |
           (
             biome == "marine biome (ENVO_00000447)"  &
               material == "sediment (ENVO_00002007)"
           ) |
           (
             biome == "marine biome (ENVO_00000447)"  &
               material == "microbial mat (ENVO_01000008)"
           ) |
           (biome == "marine coral reef biome (ENVO_01000049)") |
           (biome == "marine hydrothermal vent biome (ENVO_01000030)") |
           (biome == "marine neritic benthic zone biome (ENVO_01000025)") |
           (
             biome == "marine pelagic biome (ENVO_01000023)"  &
               material == "marine sediment (ENVO_03000033)"
           )
  ) %>%
  filter(material !=  "sea water (ENVO_00002149)")


terrestrial <- metadata_table %>%
  filter((biome == 'cropland biome (ENVO_01000245)') |
```

```
        (
          biome == 'forest biome (ENVO_01000174)'  &
            material == 'forest soil (ENVO_00002261)'
        ) |
        (
          biome == 'forest biome (ENVO_01000174)'  &
            material == 'soil (ENVO_00001998)'
        ) |
        (
          biome == 'forest biome (ENVO_01000174)'  &
            material == 'fecal material (ENVO_00002003)'
        ) |
        (
          biome == 'forest biome (ENVO_01000174)'  &
            material == 'sediment (ENVO_00002007)'
        ) |
        (
          biome == 'forest biome (ENVO_01000174)'  &
            material == 'plant litter (ENVO_01000628)'
        ) |
        (
          biome == 'grassland biome (ENVO_01000177)'  &
            material == 'organismal entity (PCO_0000031)'
        ) |
        (biome == 'terrestrial biome (ENVO_00000446)') |
        (
          biome == 'freshwater biome (ENVO_00000873)'  &
            material == 'industrial waste material (ENVO_00002267)'
        ) |
        (
          biome == 'freshwater biome (ENVO_00000873)' &
            material == 'sediment (ENVO_00002007)'
        ) |
        (
          biome == 'temperate broadleaf forest biome (ENVO_01000202)'  &
            material == 'forest soil (ENVO_00002261)'
        ) |
        (
          biome == 'temperate coniferous forest biome (ENVO_01000211)'  &
            material == 'forest soil (ENVO_00002261)'
        ) |
        (
          biome == 'tropical broadleaf forest biome (ENVO_01000200)'  &
            material == 'forest soil (ENVO_00002261)'
        ) |
        (
          biome == 'woodland biome (ENVO_01000208)'  &
            material == 'sediment (ENVO_00002007)'
        ) |
        (
          biome == 'woodland biome (ENVO_01000208)'  &
            material == 'soil (ENVO_00001998)'
        )
```

```
  )


hypersaline <- metadata_table %>%
  filter((
    biome == 'grassland biome (ENVO_01000177)'  &
      material == 'hypersaline water (ENVO_00002012)'
  ) |
    (
      biome == 'aquatic biome (ENVO_00002030)' &
        material == 'saline water (ENVO_00002010)'
    ) |
    (
      biome == 'aquatic biome (ENVO_00002030)' &
        feature == 'saline lake (ENVO_00000019)'
    )
  )


freshwater <- metadata_table %>%
  filter ((
    biome == "freshwater lake biome (ENVO_01000252)"  &
      material == "industrial waste material (ENVO_00002267)"
  ) |
    (
      biome == "freshwater lake biome (ENVO_01000252)"  &
        material == "fresh water (ENVO_00002011)"
    ) |
    (
      biome == "aquatic biome (ENVO_00002030)"  &
        material == "sediment (ENVO_00002007)"
    ) |
    (
      biome == "freshwater lake biome (ENVO_01000252)" &
        material == "part of plant (FOODON_03420174)"
    ) |
    (
      biome == "freshwater lake biome (ENVO_01000252)"  &
        material == "sediment (ENVO_00002007)"
    ) |
    (biome == "freshwater lake biome (ENVO_01000252)") |
    (
      biome == "grassland biome (ENVO_01000177)"  &
        material == "fresh water (ENVO_00002011)"
    ) |
    (
      biome == "grassland biome (ENVO_01000177)" &
        material == "sediment (ENVO_00002007)"
    ) |
    (
      biome == "grassland biome (ENVO_01000177)"  &
        material == 'brackish water (ENVO_00002019)'
    ) |
```

```
    (
      biome == "freshwater biome (ENVO_00000873)"  &
        material == "fresh water (ENVO_00002011)"
    ) |
    (
      biome == "small river biome (ENVO_00000890)"  &
        material == "fresh water (ENVO_00002011)"
    ) |
    (
      biome == "aquatic biome (ENVO_00002030)"  &
        material == "biofilm material (ENVO_01000156)"
    ) |
    (
      biome == "aquatic biome (ENVO_00002030)"  &
        material == "water (ENVO_00002006)"
    ) |
    (
      biome == "aquatic biome (ENVO_00002030)"  &
        material == "industrial waste material (ENVO_00002267)"
    ) |
    (
      biome == "freshwater river biome (ENVO_01000253)"  &
        material == "fresh water (ENVO_00002011)"
    ) |
    (
      biome == "urban biome (ENVO_01000249)"  &
        material == "sewage (ENVO_00002018)"
    )
  )  %>% filter(feature != "saline lake (ENVO_00000019)")
```

```
reference_data <- metadata_table

new_data <-
  bind_rows(hypersaline,
            terrestrial,
            marine_benthos,
            marine_pelagic,
            freshwater)

missing_ones <- setdiff(reference_data, new_data)
```

```
#add the missing samples to the biome_tables

marine_benthos <- marine_benthos %>%
  add_row(filter(
    metadata_table,
    raw_env %in% c(
      'Deep benthic microbial mat',
      'Shallow benthic microbial mat',
      'Littoral microbial mat'
    )
  ))
```

```r
terrestrial <- terrestrial %>%
  add_row(filter(
    metadata_table,
    raw_env %in%  c('snow pack',
                    'plateau',
                    'Agricultural field')
  ))

freshwater <- freshwater %>%
  add_row(filter(
    metadata_table,
    raw_env %in% c(
      'lake epilimnion',
      'lake hypolimnion',
      'water track',
      'lake outlet'
    )
  ))

marine_pelagic <- marine_pelagic %>%
  add_row(filter(metadata_table, raw_env == "mixing zone Arctic Ocean"))
```

```r
#exchange samples that were filtered incorrectly

marine_pelagic <- marine_pelagic %>%
  add_row(filter(
    freshwater,
    sample %in% c('SRR2799342',
                  'SRR2799337',
                  'SRR2799334')
  ))

marine_pelagic <- marine_pelagic %>%
  add_row(filter(metadata_table, project %in% c("PRJEB9943", "UniEuk_StoeckFilkerForster") ))

freshwater <- filter(freshwater, !sample %in% c('SRR2799342',
                        'SRR2799337',
                        'SRR2799334'))

marine_benthos <- marine_benthos %>%
  add_row(filter(terrestrial, sample %in% c('ERR2733069')))

terrestrial<- filter(terrestrial, !sample %in% c('ERR2733069'))
```

```r
#check again for missing samples

new_data <- bind_rows(hypersaline, terrestrial, marine_benthos, marine_pelagic, freshwater)

missing_ones <- setdiff(reference_data, new_data)

freshwater <- freshwater %>%
  add_row(filter(missing_ones, envplot %in% c("land_freshwater", "land_water", "land_org", "land_sedimen
```

```r
#check again for missing
new_data <- bind_rows(hypersaline, terrestrial, marine_benthos, marine_pelagic, freshwater)

missing_ones <- setdiff(reference_data, new_data)
```

##### Add Environmental information and combine datasets

```r
marine_pelagic <- marine_pelagic %>%
  add_column(environment="marine\npelagic")

marine_benthos <- marine_benthos %>%
  add_column(environment="marine\nbenthos")

terrestrial <- terrestrial %>%
  add_column(environment="terrestrial")

freshwater <- freshwater %>%
  add_column(environment="freshwater")

hypersaline <- hypersaline %>%
  add_column(environment="hypersaline")


meta_all_samples <-
  rbind(marine_pelagic,
        marine_benthos,
        terrestrial,
        freshwater,
        hypersaline)
```

##### Clean dataset from all 'canonical' microsporidia

##### Check and join datasets

```r
semi_join(otu_table, taxonomy_microsporidia, by = "amplicon") %>%
  pivot_longer(cols = -amplicon,
               names_to = "samples",
               values_to = "reads") %>%
  count(samples, wt = reads, name = "reads") %>%
  filter(reads > 1) %>%
  pull(samples) -> good_samples


otu_table %>%
  semi_join(taxonomy_microsporidia, by = "amplicon") %>%
  select(amplicon, any_of(good_samples)) %>%
  pivot_longer(cols = -amplicon,
               names_to = "sample",
               values_to = "reads") %>%
```

```
  pivot_wider(names_from = amplicon, values_from = reads) %>%
  semi_join(meta_all_samples, by = "sample") -> OTU_microsporidia

meta_all_samples %>%
  semi_join(OTU_microsporidia, by = "sample") %>%
  column_to_rownames(var = "sample") -> meta_microsporidia

OTU_microsporidia %>%
  column_to_rownames(var = "sample") %>%
  t() -> OTU_microsporidia_matrix

column_to_rownames(taxonomy_microsporidia, "amplicon") -> taxonomy_microsporidia
```

```
save(OTU_microsporidia, meta_microsporidia, taxonomy_microsporidia, file = "EukBank_microsporida_cleaned
write.table(meta_microsporidia, "META_EukBank_microsporidia_cleaned.metadata")
write.table(OTU_microsporidia, "ASV_EukBank_microsporidia_cleaned.table")
write.table(taxonomy_microsporidia, "Taxo_EukBank_microsporidia_cleaned.taxo")
```

#### Distribution of samples with and without short-branch microsporidan OTU's

```
c("#3399CC", "#CC6699", "#FFCC00", "#993333", "#669900") -> colours


world_map <- map_data("world")


ggplot(world_map, aes(x = long, y = lat, group = group)) +
  geom_polygon(fill = "grey90",
               colour = "grey90",
               size = 0.2) +
  theme_light() +
  theme(panel.background = element_rect(fill = "white"),
        panel.grid = element_blank()) -> world_map_layout


world_map_layout +
  geom_point(
    data = meta_all_samples %>% filter(!is.na(longitude)),
    aes(longitude, latitude, color = environment),
    size = 1.4,
    inherit.aes = F
  ) +
  scale_color_manual(values = colours) +
  scale_shape_manual(values = 7) +
  ggtitle("all samples") +
  guides(colour = guide_legend(override.aes = list(size = 4))) +
  theme(
    plot.title = element_text(
      size = 14,
```

```
      hjust = 0.5,
      vjust = 2
    ),
    legend.text = element_text(size = 12),
    legend.title = element_blank(),
    axis.text =    element_blank(),
    legend.position = "none"
  ) +
  xlab("") +
  ylab("") -> map_all_samples


world_map_layout +
  geom_point(
    data = meta_microsporidia %>% filter(!is.na(longitude)),
    aes(longitude, latitude, color = environment),
    size = 1.4,
    inherit.aes = F
  ) +
  scale_color_manual(values = colours) +
  scale_shape_manual(values = 7) +
  ggtitle("samples with short-branch microsporidia") +
  guides(colour = guide_legend(override.aes = list(size = 4))) +
  theme(
    plot.title = element_text(
      size = 14,
      hjust = 0.5,
      vjust = 2
    ),
    legend.text = element_text(size = 12, colour = "black"),
    legend.title = element_text(
      face = "bold",
      colour = "black",
      size = 13
    ),
    axis.text =    element_blank(),
    legend.position = "bottom"
  ) +
  xlab("") +
  ylab("") -> map_microsporidia


ggarrange(
    map_all_samples,
    map_microsporidia,
    common.legend = F,
    nrow = 2,
    heights = 2
  ) -> map_combined
```

### Similarity to reference database

#### Compute abundance values

```
OTU_microsporidia %>%
  pivot_longer(cols = -sample,
               names_to = "OTU_names",
               values_to = "read_abundance") %>%
  count(OTU_names, wt = read_abundance, name = "abundance") %>%
  filter(abundance > 0) -> OTU_T


taxonomy_microsporidia %>%
  rownames_to_column(var = "OTU_names") %>%
 select(-abundance) -> a

  a %>% left_join(ASV_T, by = "OTU_names")  -> taxonomy_microsporidia_updated_abundances

tax_reads<- taxonomy_microsporidia_updated_abundances %>%
    count(similarity, wt = abundance, name = "reads") %>%
    arrange(desc(similarity))

tax_otus<- taxonomy_microsporidia_updated_abundances %>%
    count(similarity, name = "sequences") %>%
    arrange(desc(similarity))
```

#### Plot similarity values of reads and OTUs

```
list(
  geom_segment(
    aes(xend = similarity, yend = 0),
    colour = "darkred",
    size = 2
  ) ,
  xlab("similarity to reference database (%)") ,
  theme_bw() ,
  theme(
    axis.text = element_text(size = 20, colour = "black"),
    axis.title = element_text(size = 20),
    axis.title.x = element_text(hjust = (0.5), vjust = (-2)),
    axis.title.y = element_text(vjust = (10)),
    plot.margin =  unit(c(1.5, 1.5, 1.5, 3.0), "cm")
  )
) -> stampa_theme

ggplot(tax_otus, aes(x = similarity, y = sequences)) +
  stampa_theme +
  ylab("number of OTUs") -> stampa_otus

ggplot(tax_reads, aes(x = similarity, y = reads)) +
```

```r
  stampa_theme +
  ylab("number of reads") +
  scale_y_continuous(labels = comma) -> stampa_reads


ggpubr::ggarrange(
  stampa_reads,
  stampa_otus,
  common.legend = F,
  nrow = 2,
  heights = 2
) -> stampa_merge
```

#### Venn-diagramms

##### Unique and shared OTU's across the different environments

```r
environments <-
  c("freshwater",
    "hypersaline",
    "marine\nbenthos",
    "marine\npelagic",
    "terrestrial")

create_venn <- function(environment) {
  veg_terrestrial <-
    phyloseq::prune_samples(sample_data(physeq_object)$environment == environment,
                  physeq_object)

  veg_terrestrial_clean <-
    apply(phyloseq::otu_table(veg_terrestrial), 1, function(x) {
      sum(x) > 0
    })

  veg_terrestrial_sub <-
    phyloseq::prune_taxa(veg_terrestrial_clean, veg_terrestrial)

  veg_terrestrial_asvs <-
    rownames(phyloseq::otu_table(veg_terrestrial_sub))
}


veg_list <-
  list(
    "freshwater" = create_venn("freshwater"),
    "hypersaline" = create_venn("hypersaline"),
    "marine benthos" = create_venn("marine\nbenthos"),
    "marine pelagic" = create_venn("marine\npelagic"),
    "terrestrial" = create_venn("terrestrial")
  )
```

```
venn::venn(
  veg_list,
  ilabels = "counts",
  zcolor = colours,
  borders = T,
  ellipse = T,
  ggplot = T,
  par = T,
  opacity = 0.6,
  ilcs = 2,
  sncs = 2,
) -> venn_diagram
```

**Shared ASV's**

```
Reduce(intersect,
   list(
    create_venn("freshwater"),
    create_venn("hypersaline"),
    create_venn("marine\nbenthos"),
    create_venn("marine\npelagic"),
    create_venn("terrestrial")
  )
)
```

#### Richness across different environments

```
#transform into phyloseq objects

OTU_phyloseq = otu_table(OTU_microsporidia_matrix, taxa_are_rows = T)
meta_phyloseq = sample_data(meta_microsporidia)
taxo_phyloseq = tax_table(as.matrix(taxonomy_microsporidia))


merge_phyloseq(OTU_phyloseq, meta_phyloseq, taxo_phyloseq) -> physeq_object


#output total otu abundance
summary(sample_sums(physeq_object))


#confim you don't have any empty samples
physeq_object <- phyloseq::prune_taxa(taxa_sums(physeq_object) > 1, physeq_object)
```

```
rarecurve <- vegan::rarecurve(label = F, t(otu_table(physeq_object)), step=50, cex=0.5)
```

#### Calculate richness

```
data.frame(
  "Observed" = phyloseq::estimate_richness(physeq_object, measures = "Observed"),
  "Chao1" = phyloseq::estimate_richness(physeq_object, measures = "Chao1"),
  "Environment" = phyloseq::sample_data(physeq_object)$environment
) %>%
  rename(observed = "Observed", estimated = "Chao1.Chao1") %>%
  select(-Chao1.se.chao1) -> adiv
```

```
c("freshwater",
    "hypersaline",
    "marine\nbenthos",
    "marine\npelagic",
    "terrestrial") %>%
  rep(times = 2) -> environments

c("estimated", "observed") %>% rep(times = 5) -> richnesses

run_shapiro <- function(environment, richness) {
  adiv %>%
    filter(Environment == environment) %>%
    rstatix::shapiro_test(. , vars = richness)
}


map2_dfr(environments, richnesses, run_shapiro) %>%
  mutate(environments = environments) -> shapiro_richness


rm(environments, richnesses, run_shapiro)
```

```
histogram(~observed|Environment, data = adiv) -> histogram_environments
```

#### Statistics

```
adiv %>%
group_by(Environment) %>%
summarize(Mean = median(observed, na.rm=TRUE)) -> median_richness_observed


adiv %>%
group_by(Environment) %>%
summarize(Mean = median(estimated, na.rm=TRUE)) -> median_richness_estimated

adiv %>% rstatix::wilcox_test(estimated~Environment) -> wilcox_output

saveRDS(physeq_object, "phyloseq_object_microsporidia")

ab<- readRDS("phyloseq_object_microsporidia")
```

```
write.csv(y, file = "wilcox-test-alpha-div.csv")
```

```
## Warning in rm(median_richness_estimated, median_richness_observed): object
## 'median_richness_estimated' not found

## Warning in rm(median_richness_estimated, median_richness_observed): object
## 'median_richness_observed' not found
```

##### Plot richness

```
list(
  geom_jitter(aes(color = Environment), size = 2),
    geom_boxplot(aes(color = NULL), alpha = 0.1, outlier.color = NA),
    scale_color_manual(values = colours),
   scale_shape_manual(values = 7),
    theme(
      panel.grid.major.x = element_line(size = 0.5, colour = "#f0f0f0"),
      panel.grid.major.y = element_blank() ,
      panel.grid.minor = element_blank(),
      panel.background = element_blank(),
      panel.border = element_rect(
        colour = "black",
        fill = NA,
        size = 1
      ),
      axis.line = element_line(colour = "black"),
      legend.position = ("none"),
      axis.text = element_text(colour = "black", size = 14),
      strip.placement = "outside",
      strip.text = element_text(size = 16),
      axis.title.x = element_text(size = 18, vjust = -2),
      axis.title.y = element_text(
        size = 18,
        angle = 90,
        vjust = 5
      ),
      plot.margin = unit(c(1, 0.5, 1, 1), "cm")
    )
  ) -> my_theme


## use in multiple plots
adiv %>%
  gather(key = metric, value = value, c("observed", "estimated")) %>%
  mutate(metric = factor(metric, levels = c("observed", "estimated"))) %>%
  ggplot(aes(x = Environment, y = value)) +
  coord_flip() +
  xlab("environment") +
  ylab("number of OTUs") +
  facet_grid(~ metric, scales = "free") +
  my_theme -> alpha_plot
```

##### Most abundant samples

```
meta_microsporidia %>%
  rownames_to_column(var = "sample") -> meta

adiv %>% rownames_to_column(var = "sample") %>%
  filter(observed > 100) %>%
  select(sample, observed) %>%
  left_join(meta, by = "sample") -> samples_max_OTU
```

#### OTU-heterogeneity across environments

```
presab <- phyloseq_standardize_otu_abundance(physeq_object, method = "pa")

nmds.jaccard <- ordinate(presab, method="NMDS", distance="jaccard")
```

##### NMDS (jaccard)

```
phyloseq::plot_ordination(presab, nmds.jaccard) +
  geom_jitter(aes(color = environment), size = 3) +
  scale_colour_manual(values = colours)  +
  theme_light() +
  theme(
    panel.grid = element_blank(),
    legend.title = element_text(face = "bold", size = 16),
    legend.text = element_text(size = 14),
    legend.position = c(0.905, 0.15),
    legend.background = element_rect(fill = "white", color = "black"),
    axis.line = element_line(colour = "black"),
    axis.text = element_text(size = 16),
    axis.title = element_text(size = 17),
    panel.border = element_rect(
      colour = "black",
      fill = NA,
      size = 1
    ),
    plot.margin = unit(c(1, 1, 1, 1), "cm")
  ) +
  labs(colour = "environment")  -> nmds_jaccard

nmds_jaccard + annotate("text", x=2.5, y=1.4, label="stress = 0.015") -> nmds_jaccard

ggsave("nmds.jpeg",nmds_jaccard, height=9, width=11)
```

#### Session Info

```r
sessionInfo()
```

```
## R version 4.3.1 (2023-06-16)
## Platform: aarch64-apple-darwin20 (64-bit)
## Running under: macOS Sonoma 14.3.1
##
## Matrix products: default
## BLAS:   /Library/Frameworks/R.framework/Versions/4.3-arm64/Resources/lib/libRblas.0.dylib
## LAPACK: /Library/Frameworks/R.framework/Versions/4.3-arm64/Resources/lib/libRlapack.dylib;  LAPACK v
##
## locale:
## [1] en_US.UTF-8/en_US.UTF-8/en_US.UTF-8/C/en_US.UTF-8/en_US.UTF-8
##
## time zone: Europe/Berlin
## tzcode source: internal
##
## attached base packages:
## [1] stats     graphics  grDevices utils     datasets  methods   base
##
## other attached packages:
##  [1] scales_1.3.0    venn_1.12       remotes_2.4.2.1 rstatix_0.7.2
##  [5] ggthemes_5.0.0  ggmap_4.0.0     maps_3.4.2      vegan_2.6-4
##  [9] lattice_0.22-5  permute_0.9-7   ggpubr_0.6.0    metagMisc_0.5.0
## [13] phyloseq_1.46.0 lubridate_1.9.3 forcats_1.0.0   stringr_1.5.1
## [17] dplyr_1.1.4     purrr_1.0.2     readr_2.1.5     tidyr_1.3.0
## [21] tibble_3.2.1    ggplot2_3.4.4   tidyverse_2.0.0
##
## loaded via a namespace (and not attached):
##  [1] ade4_1.7-22             tidyselect_1.2.0        Biostrings_2.70.1
##  [4] bitops_1.0-7            fastmap_1.1.1           RCurl_1.98-1.14
##  [7] digest_0.6.34           timechange_0.3.0        lifecycle_1.0.4
## [10] cluster_2.1.6           survival_3.5-7          magrittr_2.0.3
## [13] compiler_4.3.1          rlang_1.1.3             tools_4.3.1
## [16] igraph_1.6.0            utf8_1.2.4              yaml_2.3.8
## [19] data.table_1.14.10      ggsignif_0.6.4          knitr_1.45
## [22] plyr_1.8.9              abind_1.4-5             withr_3.0.0
## [25] BiocGenerics_0.48.1     grid_4.3.1              stats4_4.3.1
## [28] fansi_1.0.6             multtest_2.58.0         biomformat_1.30.0
## [31] colorspace_2.1-0        Rhdf5lib_1.24.1         iterators_1.0.14
## [34] MASS_7.3-60.0.1         cli_3.6.2               rmarkdown_2.25
## [37] crayon_1.5.2            generics_0.1.3          rstudioapi_0.15.0
## [40] httr_1.4.7              reshape2_1.4.4          tzdb_0.4.0
## [43] ape_5.7-1               rhdf5_2.46.1            zlibbioc_1.48.0
## [46] splines_4.3.1           parallel_4.3.1          XVector_0.42.0
## [49] vctrs_0.6.5             Matrix_1.6-5            carData_3.0-5
## [52] jsonlite_1.8.8          car_3.1-2               IRanges_2.36.0
## [55] hms_1.1.3               S4Vectors_0.40.2        jpeg_0.1-10
## [58] foreach_1.5.2           glue_1.7.0              admisc_0.34
## [61] codetools_0.2-19        stringi_1.8.3           gtable_0.3.4
## [64] GenomeInfoDb_1.38.5     munsell_0.5.0           pillar_1.9.0
```

```
## [67] htmltools_0.5.7         rhdf5filters_1.14.1     GenomeInfoDbData_1.2.11
## [70] R6_2.5.1                evaluate_0.23           Biobase_2.62.0
## [73] png_0.1-8               backports_1.4.1         broom_1.0.5
## [76] Rcpp_1.0.12             nlme_3.1-164            mgcv_1.9-1
## [79] xfun_0.41               pkgconfig_2.0.3
```

```r
R.Version()
```

```
## $platform
## [1] "aarch64-apple-darwin20"
##
## $arch
## [1] "aarch64"
##
## $os
## [1] "darwin20"
##
## $system
## [1] "aarch64, darwin20"
##
## $status
## [1] ""
##
## $major
## [1] "4"
##
## $minor
## [1] "3.1"
##
## $year
## [1] "2023"
##
## $month
## [1] "06"
##
## $day
## [1] "16"
##
## $`svn rev`
## [1] "84548"
##
## $language
## [1] "R"
##
## $version.string
## [1] "R version 4.3.1 (2023-06-16)"
##
## $nickname
## [1] "Beagle Scouts"
```

```r
knitr::write_bib(c(.packages()), "packages.bib")
```

```
## tweaking maps
```

```r
rm(list = ls())
```
