## Supplementary material for "O Short-branch Microsporidia, Where Art Thou? Identifying diversity hotspots for future sampling": Wilcoxon rank sum test for pairwise differences in richness among the environments

Supplementary Table 1: Wilcoxon rank sum test for pairwise differences in richness among the environments.

| group1 | group2 | n1 | n2 | statistic | p.adj | p.adj.signif |
|---|---|---|---|---|---|---|
| **Freshwater** | Hypersaline | 374 | 6 | 1989.5 | 0.006 | ** |
| **Freshwater** | Marine benthos | 374 | 435 | 124520 | 1.68E-39 | **** |
| **Freshwater** | Marine pelagic | 374 | 830 | 241347 | 1.22E-56 | **** |
| **Freshwater** | Terrestrial | 374 | 1818 | 322390 | 0.46 | ns |
| **Hypersaline** | Marine benthos | 6 | 435 | 861.5 | 0.46 | ns |
| **Hypersaline** | Marine pelagic | 6 | 830 | 1724.5 | 0.46 | ns |
| **Hypersaline** | Terrestrial | 6 | 1818 | 891.5 | 0.002 | ** |
| **Marine benthos** | Marine pelagic | 435 | 830 | 187477.5 | 0.46 | ns |
| **Marine benthos** | Terrestrial | 435 | 1818 | 152239.5 | 3.60E-88 | **** |
| **Marine pelagic** | Terrestrial | 830 | 1818 | 279632 | 2.92E-150 | **** |
